# IL-10 and ICOS differentially regulate T cell responses in the brain during chronic *Toxoplasma gondii* infection

**DOI:** 10.1101/418665

**Authors:** Carleigh A. O’Brien, Samantha J. Batista, Katherine M. Still, Tajie H. Harris

## Abstract

Control of chronic CNS infection with the parasite *Toxoplasma gondii* requires an ongoing T cell response in the brain. Immunosuppressive cytokines are also important for preventing lethal immunopathology during chronic infection. To explore the loss of suppressive cytokine exclusively during the chronic phase of infection we blocked IL-10 receptor (IL-10R). Blockade was associated with widespread changes in the inflammatory response, including increased antigen presenting cell (APC) activation, expansion of CD4+ T cells, and increased neutrophil recruitment to the brain, consistent with previous reports. We then sought to identify regulatory mechanisms contributing to IL-10 production, focusing on ICOS (inducible T cell costimulator), a molecule that promotes IL-10 production in many systems. Unexpectedly, ICOS-ligand (ICOSL) blockade led to a local expansion of effector T cells in the inflamed brain without affecting IL-10 production or APC activation. Instead, we found that ICOSL blockade led to changes in T cells associated with their proliferation and survival. Specifically, we observed increased expression of IL-2 associated signaling molecules, including CD25, STAT5 phosphorylation, Ki67, and Bcl-2 in T cells in the brain. Interestingly, increases in CD25 and Bcl-2 were not observed following IL-10R blockade. Also unlike IL-10R blockade, ICOSL blockade led to an expansion of both CD8+ and CD4+ T cells in the brain, with no expansion of peripheral T cell populations or neutrophil recruitment to the brain Overall, these results suggest that IL-10 and ICOS differentially regulate T cell responses in the brain during chronic *T. gondii* infection.

## Introduction

Immune responses have intricately evolved to protect hosts from a wide range of potentially harmful pathogens [1, 2], yet these same inflammatory responses can often cause host damage themselves. The importance of a balanced immune response is apparent in models of infection, where inflammation is required for pathogen control and survival, yet amplified immune responses observed after depletion of regulatory T cells or immunosuppressive cytokines often leads to exacerbated tissue pathology and increased mortality [3-8]. One such immunosuppressive cytokine, IL-10, has been broadly studied in the context of both tissue homeostasis and during infection, and has been shown to play a key role in suppressing many aspects of an immune response. Production of IL-10 during immune responses to infection has been attributed to a wide variety of cell types, including T cells, dendritic cells, macrophages, NK cells, and B cells [9]. IL-10 also acts on a wide range of cell types, with one of its main roles being the downregulation of MHC and costimulatory molecules in antigen presenting cells (APCs), thereby preventing their full activation capacity and limiting T cell responses [10-12]. IL-10 also has direct effects on T cells, limiting IFNγ and IL-2 production, as well as T cell proliferation *in vitro* [13, 14].

Infection with the eukaryotic parasite *Toxoplasma gondii* leads to widespread activation of the immune system and systemic inflammation that is required for host survival [15]. The generation of a parasite-specific adaptive immune response clears the parasite from most peripheral tissues; however, the parasite is able to encyst in the central nervous system and establish a chronic infection [16, 17]. This chronic infection requires ongoing activation and infiltration of highly polarized Th1 cells into the brain in order to prevent extensive parasite replication and fatal disease [18, 19]. However, like in other models of infection, regulation of this immune response is also required to promote host survival. In particular, IL-10 production is required for host survival during acute infection. IL-10 knockout mice succumb to CD4+ T cell-mediated immunopathology and excessive inflammatory cytokine production in the periphery early in the course of infection [3, 20]. Similarly, continued IL-10 production in the chronic phase of infection is also necessary for host survival. IL-10 knockout mice given the antibiotic sulfadiazine early in the infection to limit parasite replication survive the acute phase of infection, but later present with similar CD4+ T cell-mediated fatal immunopathology in the brain [21]. Despite demonstrating the requirement for IL-10 signaling over the course of *T. gondii* infection, previous studies have not addressed what additional signals promote immune regulation in the context of chronic neuroinflammation.

ICOS (inducible T cell costimulator) is a costimulatory molecule expressed on activated T cells [22, 23]. ICOS signaling is important in a wide variety of immune responses, including optimal antibody class switching and T cell inflammatory cytokine production [23-26]. The primary function attributed to ICOS is the amplification of effector T cell responses by serving as a costimulatory molecule similar to its family member CD28 [27]. More recently, ICOS has been shown to also promote immune regulation through potent induction of IL-10 both *in vitro* and in mouse models of acute inflammation [28-30]. In addition to promoting production of IL-10, ICOS is also important for maintaining effector regulatory T cell populations. During homeostasis, blockade of ICOS signaling results in a loss of CD44^hi^CD62L^lo^ effector T_regs_ in the spleen [31, 32]. In mouse models of diabetes and helminth infection, a similar loss of Tregs is seen with a lack of ICOS signaling, in addition to decreased IL-10 production [33, 34]. During acute *T. gondii* infection, ICOS signaling has been reported to amplify T cell inflammatory responses by promoting increased IFNγ production early in the infection [35, 36]. ICOS appears to play a redundant role with CD28 in this setting, as mice lacking ICOS are only more susceptible to infection on a CD28-/-background [35]. Together, these reports highlight the context-dependent role of ICOS signaling, contingent on variables ranging from the type of inflammatory environment to stage of infection [22-36].

The role of ICOS, and its relationship to IL-10–mediated regulation of immune responses, in the context of a chronic neuroinflammatory response to a pathogen is not well understood. In this study, we first characterized what role IL-10 plays in promoting regulation of immune responses in the chronic stage of infection by blocking signaling through the IL-10 receptor. This IL-10R blockade during chronic infection led to an expansion of CD4+ effector T cells correlating with increased expression of CD80 on APCs, along with widespread immunopathology. Based on previous reports implicating a role for ICOS in stimulating IL-10 production, we then addressed the question of whether ICOS signaling can promote suppression of chronic T cell responses in the central nervous system through induction of IL-10. Surprisingly, we find that blockade of ICOS signaling during chronic infection with *T. gondii* does not lead to decreased IL-10 production from either regulatory or effector T cells, nor does it lead to impaired inflammatory cytokine production. In fact, despite maintaining IL-10 levels in the brain, blockade of ICOSL still results in a loss of T cell regulation, with two to three-fold more effector T cells found in the inflamed brain. Interestingly, this increase in effector T cells occurred without a loss of T_regs_ in the brain and did not affect parasite burdens. We found this increase in T cell number in the brain to correlate with increased levels of CD25 and pSTAT5 expression in effector T cells following ICOSL blockade, suggesting increased responses to IL-2. Along these lines, ICOSL blockade increased T cell proliferation and expression of the survival factor Bcl-2 among the effector CD4+ and CD8+ T cell populations in the brain. Interestingly, IL-10R blockade did not result in the same increases in IL-2-associated signaling molecules CD25 and Bcl-2; rather, IL-10R blockade increased the activation state of APCs. Taken together, our results suggest that ICOS signaling on T cells can suppress STAT5-induced survival signals, providing a mechanism of local suppression in the context of chronic inflammation in the brain that is distinct from IL-10-mediated regulation.

## Materials and Methods

### Mice and infection model

C57Bl/6 and B6.129S6-*Il10*^*tm1Flv*^*/J* (*Tiger*) mice were purchased from Jackson Laboratories. All animals were kept in UVA specific pathogen-free facilities, and were age and sex matched for all experiments. All experimental procedures followed the regulations of the Institutional Animal Care and Use Committee at the University of Virginia. Infections used the avirulent type II *Toxoplasma gondii* strain Me49, which were maintained in chronically infected Swiss Webster mice (Charles River) and passaged through CBA/J mice (Jackson Laboratories) before experimental infections. For experimental infections, the brains of chronically infected (4 to 8 weeks) CBA/J mice were homogenized to isolate tissue cysts. Experimental mice were then injected intraperitoneally with 10 to 20 Me49 cysts.

### Antibody blockade treatments

For IL-10R blockade studies, chronically infected mice (28-35 days post-infection) were treated intraperitoneally with 200 µg of a monoclonal antibody blocking the IL-10R (BioXCell) or a control rat IgG antibody. Treatments were given every 3 days, and mice were euthanized when neurological symptoms developed between 7-10 days post-antibody treatment. For ICOSL blockade studies, chronically infected mice (28-35 days post-infection) were treated intraperitoneally with 150 µg of an α-ICOSL (CD275) blocking antibody (BioXCell) or a control rat IgG antibody. Antibody treatments were given every 3 days for 14 days, totaling 5 treatments.

### Tissue and blood processing

Mice were sacrificed and perfused with 40 mL 1X PBS, and perfused brains, spleens, and lymph nodes (pooled deep and superficial cervical) were put into cold complete RPMI media (cRPMI) (10% fetal bovine serum, 1% NEAA, 1% pen/strep, 1% sodium pyruvate, 0.1% β-mercaptoethanol). Brains were then minced with a razor blade and enzymatically digested with 0.227mg/mL collagenase/dispase and 50U/mL DNase (Roche) for 1 hour at 37°C. After enzyme digestion, brain homogenate was passed through a 70µm filter (Corning). To remove myelin, filtered brain homogenate was resuspended with 20 mL 40% percoll and spun for 25 minutes at 650g. Myelin was then aspirated and the cell pellet was washed with cRPMI, then resuspended and cells were counted.

Spleens and lymph nodes were homogenized and passed through a 40µm filter (Corning) and pelleted. Lymph node cells were then resuspended in cRPMI and counted. Spleen cells were resuspended with 2 mL RBC lysis buffer (0.16 M NH_4_Cl). Following RBC lysis, spleen cells were washed with media and then resuspended with cRPMI for counting.

For experiments in which peripheral blood was taken, mice were sacrificed and the right atrium was cut in preparation for perfusion. Before perfusion, 300µL blood was collected from the chest cavity. For isolation of circulating leukocytes, collected blood was put in 1mL heparin (100 USP/mL) to prevent clotting. Samples were then pelleted and resuspended in 2 mL RBC lysis buffer for 2 minutes. Samples were washed once with cRPMI and a second RBC lysis step was performed. Finally, blood cells were resuspended in cRPMI for staining and counting. For serum isolation, blood was allowed to clot overnight at 4°C. Samples were then spun at 14,000 rpm for 10 minutes to separate clotted blood from serum. After spinning, serum (supernatant) was transferred to a clean tube and stored at −80°C.

### ELISA

ELISAs for parasite-specific IgG were performed as previously described [37]. Briefly, Immunolon 4HBX ELISA plates (Thermofisher) were coated with 5 µg soluble *Toxoplasma* antigen (STAg) overnight at 4°C. After antigen coating, plates were washed in 1x PBS with 0.1% Triton and 0.05% Tween, then blocked with 10% FBS for 2 hours at room temperature. After washing, serial dilutions of collected serum were added to plate wells overnight at 4°C. After incubation with serum samples, plates were washed and wells were incubated with HRP (Southern Biotechnology) for 1 hour at room temperature. ABTS peroxidase substrate solution (KBL) was then added to wells and immediately after a color change plates were read on an Epoch BioTek plate reader using Gen5 2.00 software.

### Flow cytometry

Single cell suspensions from collected tissues were plated in a 96-well plate. Cells were initially incubated with 50µL Fc block (1µg/mL 2.4G2 Ab (BioXCell), 0.1% rat gamma globulin (Jackson Immunoresearch)) for 10 minutes at room temperature. Cells were then surface stained for CD3 (145-2C11), CD19 (eBio1D3), NK1.1 (PK136), ICOS (7E.17G9), ICOSL (HK5.3), MHCII (M5/114.15.2), CD25 (PC61), CD8 (S3-6.7), CD11c (N418), CD4 (GK1.5), CD80 (16-10A1), CD86 (GL1), CD45 (30-F11), CD11b (M1/70), Ly6G (1A8), and a live/dead stain for 30 minutes at 4°C. Parasite-specific cells were identified using a PE-conjugated MHCII tetramer (AVEIHRPVPGTAPPS) (National Institutes of Health Tetramer Facility). After surface staining, cells were washed with FACS buffer (1% PBS, 0.2% BSA, and 2mM EDTA) and fixed at 4°C overnight with a fixation/permeabilization kit (eBioscience) or 2% PFA. Following overnight fixation, cells were permeabilized and stained for intracellular markers Bcl-2 (3F11), Ki67 (SolA15), and Foxp3 (FJK-16S) for 30 minutes at 4°C. Cells were then washed, resuspended in FACS buffer, and run on a Gallios flow cytometer (Beckman Coulter). Analysis was done using Flowjo software, v.10.

### Cytokine restimulation

Single cell suspensions were plated into a 96-well plate. For T cell cytokine restimulation, cells were incubated at 37°C with 20 ng/mL PMA and 1 µg/mL ionomycin (Sigma) in the presence of brefeldin A (Alfa Aesar). After incubation for 5 hours, cell suspensions were washed, surface stained, and fixed. Cells were then incubated with an antibody against IFNγ (XMG1.2) for 30 minutes at 4°C. To determine IL-10 production, *Tiger* mice expressing eGFP under the IL-10 promoter were used. Cells from these mice were stimulated with PMA/ionomycin and stained for surface molecules as described above, then lightly fixed with 2% PFA for 30 minutes at room temperature. Cells were then permeabilized with permeabilization buffer (eBioscience) and incubated with a biotin-conjugated α-GFP antibody (BD Biosciences) for 30 minutes at 4°C. Cells were then washed and incubated with a PE-conjugated streptavidin secondary antibody (eBioscience) for 30 minutes at room temperature. Cells were then fixed with a fixation/permeabilization kit (eBioscience) overnight at 4°C before other intracellular staining was performed. For myeloid cell cytokine restimulation, single cell suspensions were plated into a 96-well plate and incubated with brefeldin A for 5 hours at 37°C before surface and intracellular staining for IL-12 (C17.8) (eBioscience).

### qRT-PCR

Approximately 100 mg brain tissue was put into 1 mL Trizol (Ambion) in bead beating tubes (Sarstedt) containing 1mm zirconia/silica beads (BioSpec). Using a Mini-bead beater (Biospec), tissue was homogenized for 30 seconds and RNA extraction was completed using the manufacturer’s instructions (Ambion). For cDNA synthesis, a High Capacity Reverse Transcription Kit (Applied Biosystems) was used. qRT-PCR was set up using a 2X Taq-based mastermix (Bioline) and Taqman gene expression assays (Applied Biosystems) and reactions were run on a CFX384 Real-Time System (Bio-Rad). HPRT was used as a housekeeping gene for all reactions and relative expression compared to control treated animals was calculated as 2^(-ΔΔCT)^.

### T. gondii cyst counts

After mincing with a razor blade, brain tissue was passed through an 18- and 22-gauge needle. 30µL of brain homogenate was then placed onto a microscope slide (VWR) and cysts were counted manually on a brightfield DM 2000 LED microscope (Leica).

### Immunohistochemistry

Brains from mice were embedded in OCT and flash frozen on dry ice. Samples were then stored at −20°C until cutting. For Ki67 immunofluorescence staining, fresh frozen sections were lightly fixed in 25% EtOH/75% acetone solution for 15 minutes at −20°C. Sections were then blocked in 1% PBS containing 2% goat serum (Jackson Immunoresearch), 0.1% triton, and 0.05% Tween 20 for 1 hour at room temperature, then incubated with primary antibodies at 4°C overnight. After primary staining, sections were washed with 1% PBS and incubated with secondary antibodies for 30 minutes at room temperature. Finally, sections were nuclear stained with DAPI (Thermo Scientific) for 5 minutes at room temperature. Sections were then covered with aquamount (Lerner Laboratories) and coverslips (Fisherbrand).

For pSTAT5 immunofluorescence staining, fresh frozen sections were fixed in 3.2% PFA for 20 minutes at room temperature and then permeabilized with ice cold 90% methanol for 10 minutes at −20°C [31]. Sections were then incubated with blocking buffer and an anti-pSTAT5 (D47E7) (Cell Signaling Technologies) primary antibody as described above. After primary antibody staining, pSTAT5 signal was amplified using an anti-rabbit biotinylated antibody (Jackson Immunoresearch) before using a streptavidin secondary antibody. All images were captured using a DMI6000 B widefield microscope with a Hamamatsu C10600 Orca R^2^ digital camera (Leica), and visualized using Metamorph software. Images were then analyzed using Image J software.

For quantification of the number of Ki67- or pSTAT5-expressing cells, 12-15 equal-sized pictures were taken (40x) within each brain slice, and the numbers of Ki67+ or pSTAT5+ CD4+ or CD8+ cells were counted in each image. Numbers from each picture were then averaged and reported as the average number of Ki67+ or pSTAT5+ per 100µm^2^ or 500µm^2^ per mouse, respectively.

### Statistics

Statistical analysis comparing two different groups at a single time point was performed in Prism software (v. 7.0a) using a Student’s t test. In some cases, multiple experiments from different infection dates were combined to show natural biological variation between infections. When data from multiple experiments were combined, a randomized block ANOVA was used in R v.3.4.4 statistical software. This statistical test accounted for natural variability between experimental dates by modeling the treatment (IgG vs. α-IL-10R or α-ICOSL) as a fixed effect and the experimental date as a random effect. The particular test used for each individual panel shown is specified in the figure legend. P values are displayed as follows: ns= not significant, * p<0.05, ** p<0.01, *** p<0.001. All data were graphed using Prism software, and the number of mice per group is indicated in the figure legend.

## Results

### Blockade of IL-10R during chronic *T. gondii* infection leads to broad changes in the immune response and results in fatal immunopathology in brain

During chronic infection with *T. gondii*, both effector T cells and T_regs_ recruited to the brain are capable of producing IL-10 [38]. Previously published results have shown a requirement for IL-10 signaling to limit fatal immunopathology in both the acute and chronic stages of infection [3, 21, 39, 40]. Despite the necessity for IL-10 over the course of infection with *T. gondii*, previous studies addressing the role of IL-10 in the chronic phase of infection relied on total IL-10 knockout mice, which succumb to fatal immunopathology during the first two weeks of infection [41]. In order for these mice to survive to the chronic stage of infection, the anti-parasitic drug sulfadiazine must be administered for the first two weeks of infection in order to limit parasite replication and dissemination. These previously published studies reported that, after sulfadiazine treatment, IL-10 knockout mice subsequently presented with CD4+ T cell dependent lethal immunopathology and died late in the chronic stage of disease [21]. It is still unknown, however, how a loss of IL-10 only in the chronic phase of infection influences immune responses. Thus, we treated mice with an α-IL-10R blocking antibody beginning four weeks post infection. Mice treated with an α-IL-10R blocking antibody during the chronic stage of infection presented with overt disease and became moribund between 7 to 10 days post-treatment. H&E staining of tissue sections from brains of α-IL-10R-treated mice showed increased leukocyte infiltration and associated areas of necrosis not seen in control treated animals (Figure 1A-B). The increased numbers of immune cells in the brains of α-IL-10R-treated mice included increases in both CD4+Foxp3-T cells and infiltrating macrophages (Figure 1C and Supplementary Figure 1A). Though the increase in T cell number in the brains of α-IL-10R-treated mice predominantly came from an increase in the CD4+Foxp3-T cell compartment and not the CD8+ T cell compartment, an increased frequency of both Ki67+ CD4+ and CD8+ effector T cells was observed (Figure 1D). Interestingly, using an MHCII tetramer reagent to measure CD4+ T cells specific for the parasite, we observed no significant increase in the number of tetramer-positive CD4+Foxp3-T cells (Figure 1E). Though this analysis of antigen specificity was done using only a single parasite peptide, this result suggests that IL-10R blockade in the chronic phase of infection leads to the expansion of CD4+ effector T cells with potentially different antigen specificities, possibly through the expansion of other parasite-specific T cell clones or self-antigen-specific T cell clones.

**Figure 1.**
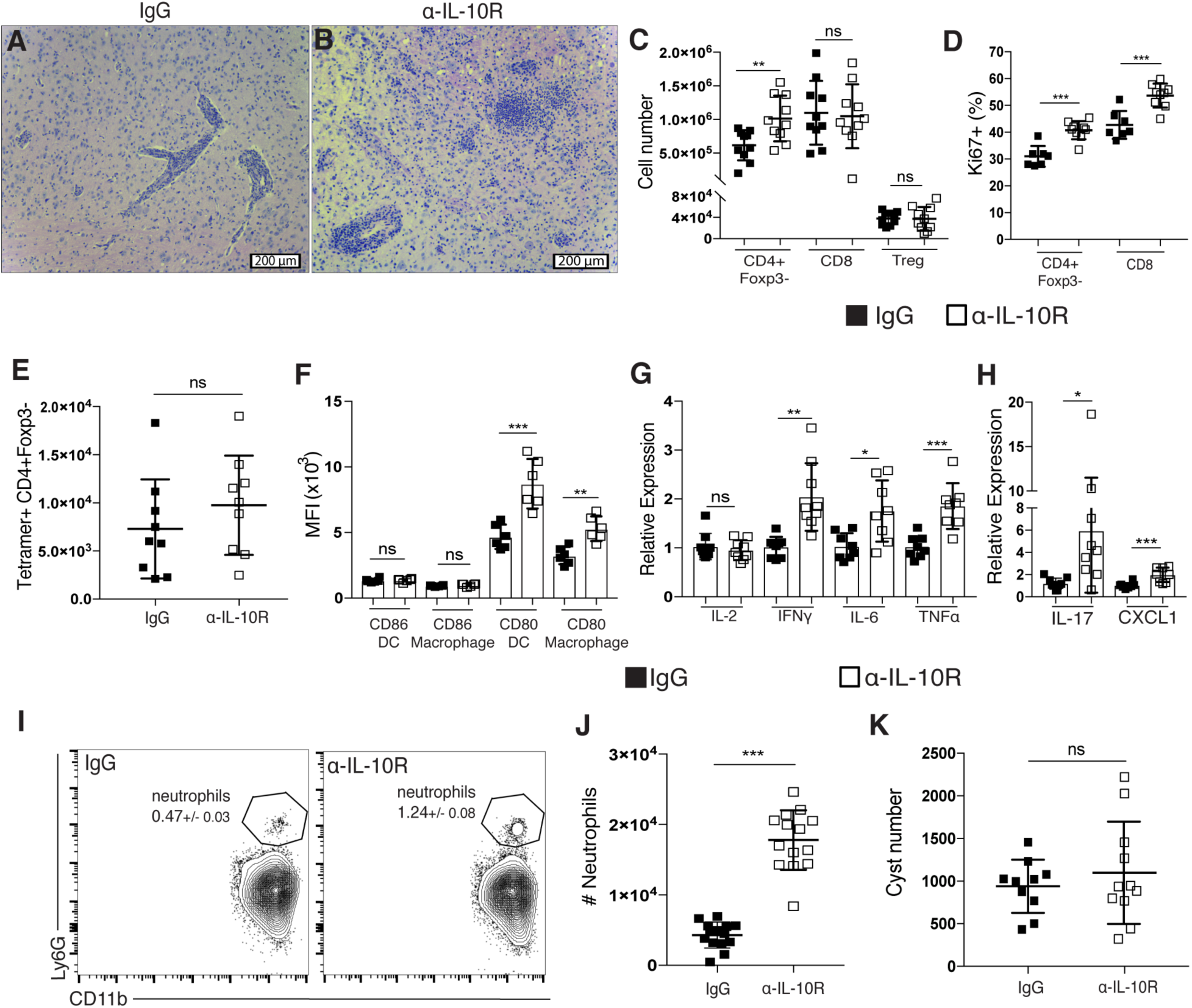
Blockade of IL-10R during chronic *T. gondii* infection leads to broad changes in the immune response and fatal immunopathology in the brain. (**A-K**) Rat IgG or an α-IL-10R blocking antibody was administered to chronically infected mice beginning at day 28 post-infection. (**A-B**) Representative H&E stained brain sections from a chronically infected control-treated mouse (**A**) and an α-IL-10R-treated mouse (**B**). (**C**) T cells isolated from the brain were analyzed by flow cytometry. Effector CD4+ T cells (CD4+Foxp3-), CD8 T cells, and T_regs_ (CD4+Foxp3+) were enumerated (n=5 per group, data is pooled from two independent experiments and analyzed using a randomized block ANOVA). (**D**) The frequency of Ki67+ effector T cells was measured by flow cytometry from mononuclear cells isolated from the brains of control and α-IL-10R-treated mice (n=3-5 per group, data is pooled from two independent experiments and analyzed using randomized block ANOVA). (**E**) Parasite-specific CD4+ effector T cells were assessed by flow cytometry using an MHCII-peptide tetramer. (**F**) The mean fluorescence intensity (MFI) of CD80 and CD86 on brain-infiltrating APCs. DCs were gated on CD45^hi^CD3^-^NK1.1^-^CD19^-^CD11c^+^MHCII^hi^ live singlet cells and macrophages were gated on CD45^hi^CD3^-^NK1.1^-^CD19^-^CD11c^-^CD11b+ live singlet cells (n=6 per group, data is representative of 3 independent experiments and analyzed using Student’s t test). (**G-H**) qRT-PCR was done using mRNA isolated from whole brains of chronically infected control or α-IL-10R-treated mice. Relative expression was normalized to the control (IgG-treated) group (n=4-5 per group, data is pooled from two independent experiments and analyzed using randomized block ANOVA). (**I**) Representative flow plots showing the neutrophil population isolated from the brains of chronically infected mice. Neutrophils were gated on CD45^hi^CD3^-^CD19^-^NK1.1^-^ CD11b^+^Ly6G^+^ live singlet cells. Number in plot indicates the mean frequency +/- standard error. (**J**) Neutrophils were identified by flow cytometry from cells isolated from the brains of chronically infected control or α-IL-10R-treated mice (n=4-5 per group, data is pooled from three independent experiments and analyzed by randomized block ANOVA). (**K**) Total cyst numbers from the brains of chronically infected control and α-IL-10R-treated mice were counted using light microscopy (n=3-4 per group, data is pooled from three independent experiments and analyzed by randomized block ANOVA). * denotes p<0.05, ** denotes p<0.01, and *** denotes p<0.001 for all panels.

The increased proliferation of effector T cells in the brain correlated with an increase in the expression of costimulatory molecule CD80 on infiltrating dendritic cells and macrophages in α-IL-10R-treated mice (Figure 1F). The increased numbers of immune cells infiltrating the brain was associated with increased mRNA levels of many pro-inflammatory cytokines and chemokines, including IFNγ, IL-6, TNFα, IL-17, and CXCL1, demonstrating a widespread increase in inflammation in the brain in the absence of IL-10 signaling (Figure 1G-H). An increase in IL-17 production is notable in this model, as infection of wild-type mice with *T. gondii* typically leads to a robust Th1-polarized immune response, characterized by IL-12 and IFNγ production that persists throughout the chronic stage, with very little production of Th17-associated cytokines [42, 43]. Indeed, production of IL-17 has been linked to a loss of IL-27-mediated regulation of immune responses during infection with *T. gondii* [43, 44], suggesting a pathogenic role for IL-17 in this context.

Production of IL-17 and CXCL1 has previously been shown to enhance recruitment of neutrophils to inflamed tissues and enhance their activity in certain disease contexts [45-50]. Neutrophil recruitment to the central nervous system however, has been shown to be detrimental in many cases [50-52]. With the increased mRNA levels of both IL-17 and CXCL1 seen in the brain after IL-10R blockade, we wanted to assess whether more neutrophils were recruited to the inflamed brain in this context. Indeed, α-IL-10R-treated mice had a nearly three-fold increase in neutrophil numbers in the brain in comparison to control-treated mice (Figure 1I-J and Supplementary Figure 1B-C). Despite the increased T cell response in the brain with IL-10R blockade, no change in the number of parasite cysts was observed (Figure 1K).

The effects of IL-10R blockade during the chronic phase of infection were not limited to the inflamed brain, as increased CD4+Foxp3-effector T cells and myeloid cells were found in the spleens of α-IL-10R-treated mice (Supplementary Figure 1D-E). Myeloid cells in the spleen were also highly activated, with higher levels of CD80 expression on APCs in the spleens of α-IL-10R-treated mice as opposed to controls (Supplementary Figure 1F). Large areas of necrosis were also observed in the livers of α-IL-10R-treated mice (Supplementary Figure 1G-H), suggesting that the increased inflammatory response seen following IL-10R blockade can contribute to extensive immunopathology in not only the brain but peripheral tissues as well. Overall, a loss of IL-10 signaling specifically during the chronic infection led to widespread immune cell activation that rapidly resulted in fatal immunopathology.

### Blockade of ICOSL does not affect IL-10 production during chronic infection, but leads to expanded T cell populations in the brain

Despite clear evidence that IL-10-mediated regulation of immune responses during *T. gondii* infection is required for host survival, little is known about what signals can induce IL-10 production in activated T cells during the chronic inflammation in the brain associated with the later stages of infection. Several studies have implicated a role for ICOS in inducing IL-10 production in cases of acute inflammation [29, 33]. We found ICOS-expressing T cells in the brain during chronic *T. gondii* infection, as well as infiltrating APCs expressing the ICOS-ligand (ICOSL) (Figure 2A-B). To address whether ICOS signaling is important for T cell production of IL-10 in the brain during chronic infection, we treated chronically infected IL-10-eGFP reporter (*Tiger*) mice with the ICOSL blocking antibody. Surprisingly, following ICOSL blockade in the chronic stage of infection, we did not see decreases in IL-10 production from either CD4+Foxp3-effector T cells or CD4+Foxp3+ regulatory T cells in the brain (Figure 2C), and IL-10 mRNA expression from whole brain homogenate was also not decreased (Figure 2D). Although levels of IL-10 were not affected after α-ICOSL treatment, we unexpectedly observed a two- to three-fold increase of both CD4+Foxp3- and CD8+ effector T cells, respectively (Figure 2E). This increase in effector T cell number was not due to a loss of the local regulatory T cell population, as their numbers in the brain were also increased, though to a lesser degree than the effector T cell populations (Figure 2E). Infiltrating myeloid cell numbers were also assessed, revealing almost a two-fold increase in dendritic cells in the brain compared to control treated animals (Supplementary Figure 2A). Though increased numbers of dendritic cells were found in the brain following α-ICOSL treatment, there was no effect on IL-12 production from either the dendritic cells or macrophages isolated from the brain (Supplementary Figure 2B-C), suggesting that continued production of IL-12 is not reliant on ICOS-ICOSL interactions. We also found an increase in the number of IFNγ-producing effector T cells in the brain, while mRNA levels of other pro-inflammatory cytokines and chemokines were not increased (Supplementary Figure 2D-E). The increased numbers of T cells in the brains α-ICOSL-treated animals could be seen dispersed throughout the brain parenchyma (Figure 2F-G), though unlike α-IL-10R treated animals, ICOSL blockade was not lethal in the observed timeframe. Additionally, we observed an increase in the number of tetramer-positive CD4+ effector T cells in the brain (Figure 2H), suggesting that ICOSL blockade can lead to an expansion of parasite-specific CD4+ effector T cells in the brain. Another distinction between IL-10R blockade and ICOSL blockade in the chronic phase of infection was a difference in neutrophil recruitment. Whereas an increase in neutrophil numbers in the brain was seen in α-IL-10R-treated mice (Figure 1), no significant increase was seen in α-ICOSL-treated mice (Figure 2I-J).

**Figure 2.**
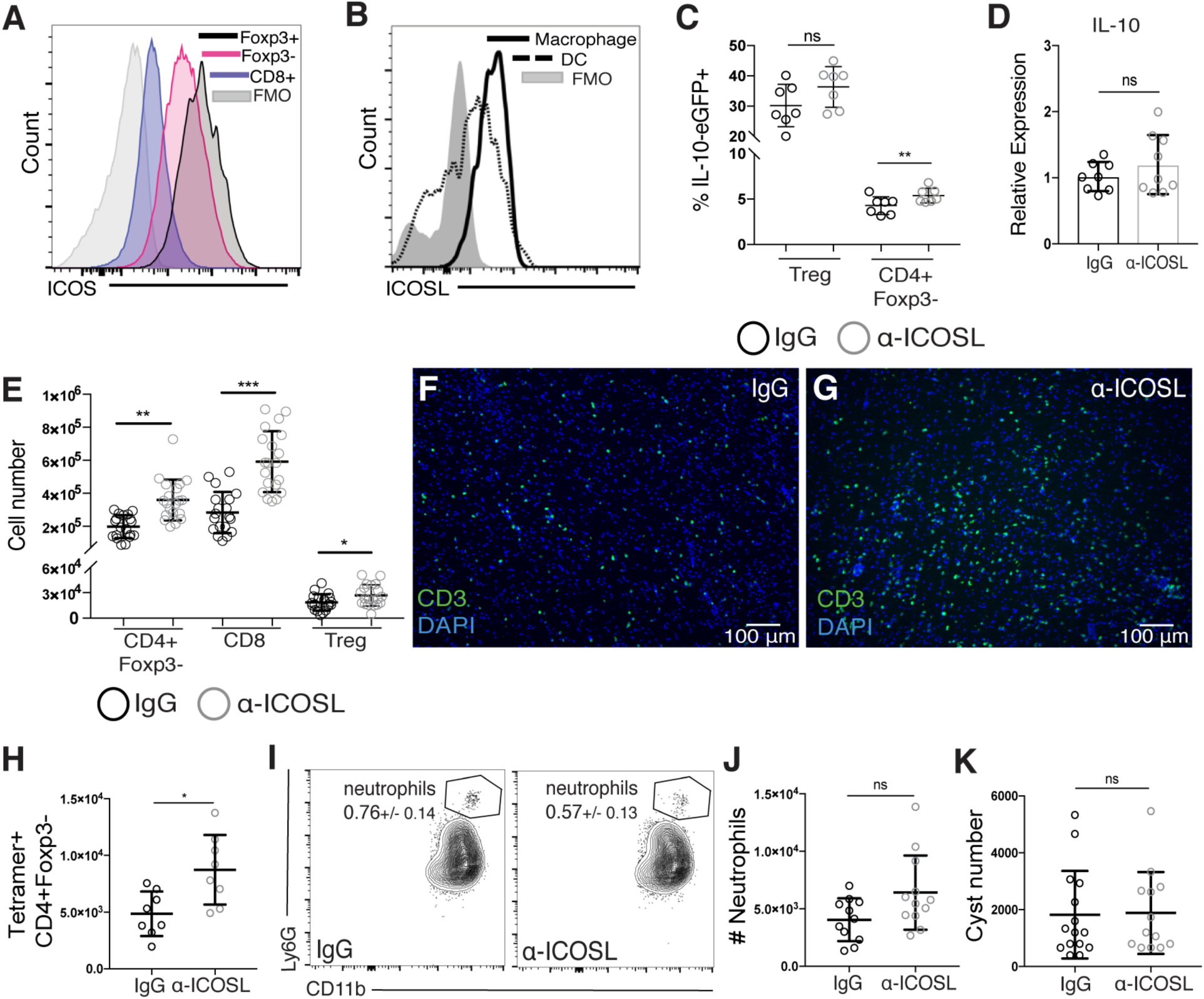
Blockade of ICOSL does not affect IL-10 production during chronic *T. gondii* infection, but leads to expanded T cell populations in the brain. (**A**) ICOS expression on T cells and (**B**) ICOSL expression on infiltrating myeloid cells isolated from the brains of chronically infected mice using flow cytometry. Immune cell populations were gated as described above. (**C**) Chronically infected IL-10-eGFP reporter (*Tiger*) mice were treated with an α-ICOSL blocking antibody beginning 4 weeks post-infection. Blocking or control antibody was given every 3 days for 2 weeks. The frequency of IL-10-GFP+ CD4 effector and T_reg_ cells isolated from the brain is shown (n=3-4 per group, data is pooled from two independent experiments and analyzed using randomized block ANOVA). (**D**) The relative expression of IL-10 mRNA in chronically infected whole brains treated with control or α-ICOSL blocking antibody. Relative expression was normalized to the control (IgG-treated) group (n=4 per group, data is pooled from two independent experiments and analyzed using randomized block ANOVA). (**E-K**) Chronically infected WT mice were treated with an α-ICOSL blocking antibody or control rat IgG. (**E**) Total T cell numbers isolated from the brain were analyzed by flow cytometry (n=3-4 per group, representative data is pooled from 5 independent experiments and analyzed by randomized block ANOVA). (**F-G**) Representative brain sections stained for CD3 (green) from control (**F**) or α-ICOSL-treated (**G**) mice. (**H**) Parasite-specific CD4+ effector T cells were assessed by flow cytometry using an MHCII-peptide tetramer. (**I**) Representative flow plots of the neutrophil population and (**J**) total numbers of neutrophils isolated from the brain. Number in plots indicates the mean frequency +/- standard error. (n=3-4 per group, data is pooled from three independent experiments and analyzed using randomized block ANOVA). (**K**) Total cyst numbers from the brains of chronically infected control and α-ICOSL-treated mice enumerated by light microscopy (n=4-5 per group, data is pooled from three independent experiments and analyzed using randomized block ANOVA). * denotes p<0.05, ** denotes p<0.01, and *** denotes p<0.001 for all panels.

Blockade of ICOSL primarily affected the T cell populations in the CNS and did not affect T cell numbers in the spleen, draining LNs, or blood of α-ICOSL-treated mice (Supplementary Figure 2-H). ICOS signaling has also been shown to be crucial for primary antibody responses to infection [26, 53, 54]. In order to assess whether the increased T cell numbers found in the brains of α-ICOSL treated animals was merely a result of a change in parasite burden due to decreased antibody production, both parasite-specific serum IgG titers and brain cyst burden was measured. No change was seen in either circulating parasite-specific IgG (Supplementary Figure 2I) or parasite burden in the brain (Figure 2K).

Together, these results suggest that ICOS signaling limits excessive T cell responses in the brain during chronic neuroinflammation independent of changes in either IL-10 production or parasite burden.

### ICOSL blockade during chronic infection is associated with increases in Ki67- and Bcl-2-expressing effector T cells in the brain

We next wanted to determine how lack of ICOS-ICOSL interaction during chronic inflammation leads to increased numbers of T cells in the CNS. Using immunofluorescence staining for Ki67, increased numbers of proliferating CD4+ and CD8+ effector T cells were found throughout the brain after ICOSL blockade during chronic infection (Figure 3A-D). Using flow cytometry, we confirmed this increase in the number of proliferating effector T cells in the brain (Figure 3E). Distinct from IL-10R blockade, the increase in Ki67+ effector T cells in the brain following ICOSL blockade was not correlated with an increase in CD80 or CD86 expression on infiltrating APCs (Figure 3F). Rather, ICOSL blockade led to the upregulation of the pro-survival factor Bcl-2 in both CD4+Foxp3- and CD8+ effector T cells isolated from the brain (Figure 3G-I). The above results suggest that ICOS limits effector T cell proliferation and survival factor expression during chronic neuroinflammation

**Figure 3.**
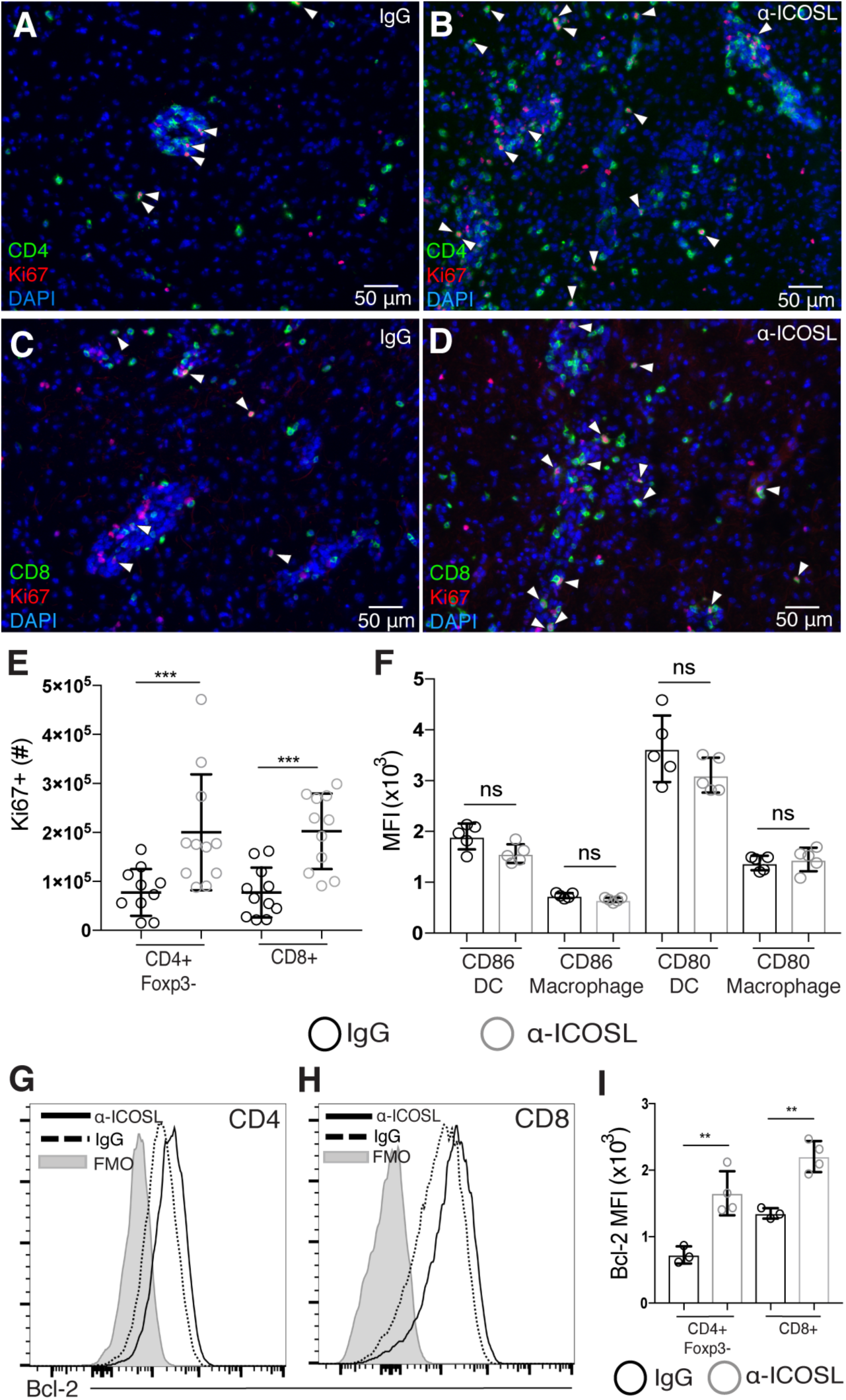
ICOSL blockade during chronic infection is associated with increases in Ki67- and Bcl-2-expressing effector T cells in the brain. (**A-D**) Brain sections from chronically infected control (**A, C**) and α-ICOSL-treated (**B, D**) mice were stained for CD4 or CD8 (green), Ki67 (red), and DAPI (blue). White arrowheads indicate Ki67+ CD4 or CD8 T cells. (**E**) The number of Ki67+ effector T cells in the brains of control or α-ICOSL-treated mice was analyzed by flow cytometry (n=3-4 mice per group, data is pooled from 3 independent experiments and analyzed by randomized block ANOVA). (**F**) The MFI of CD80 and CD86 on infiltrating APCs isolated from the brain after α-ICOSL or control treatment (n=5 mice per group, data is representative of 3 independent experiments and analyzed using Student’s t test). (**G-H**) Representative histograms of Bcl-2 expression measured by flow cytometry on effector CD4+ (**G**) and effector CD8+ (**H**) T cells isolated from the brain. (**I**) The MFI of Bcl-2 on effector T cell populations isolated from the brains of chronically infected control or α-ICOSL-treated mice (n=3-4 per group, data is representative of 4 independent experiments and analyzed using Student’s t test). * denotes p<0.05, ** denotes p<0.01, and *** denotes p<0.001 for all panels.

### Blockade of ICOSL increases CD25 expression and STAT5 phosphorylation in effector T cells in the brain during chronic infection

IL-2 signaling in T cells has been shown to support both their proliferation and survival [55, 56]. Thus, we wanted to determine if the increased effector T cell proliferation and survival factor Bcl-2 expression could be a result of increased response to IL-2 following ICOSL blockade. After ICOSL blockade, both CD4+Foxp3- and CD8+ effector T cells isolated from the brain expressed higher levels of CD25, and there was a two- to three-fold increase in the total number of CD25+ effector T cells in the brain compared to control treated animals (Figure 4A-D). In order to assess whether more of the infiltrating T cells in the brain may be responding to IL-2, we used immunofluorescence staining for phosphorylated STAT5 (pSTAT5), the main signaling molecule downstream of the IL-2R. Correlated with the increased number of CD25+ effector T cells found in the brain with ICOSL blockade, we observed increased numbers of pSTAT5-positive CD4+ and CD8+ T cells in the brains of α-ICOSL-treated animals (Figure 4E-J). Taken together, these data suggest that after ICOSL blockade, effector T cells may maintain higher levels of CD25 and pSTAT5, which in turn could support increased proliferation and Bcl-2 expression.

**Figure 4.**
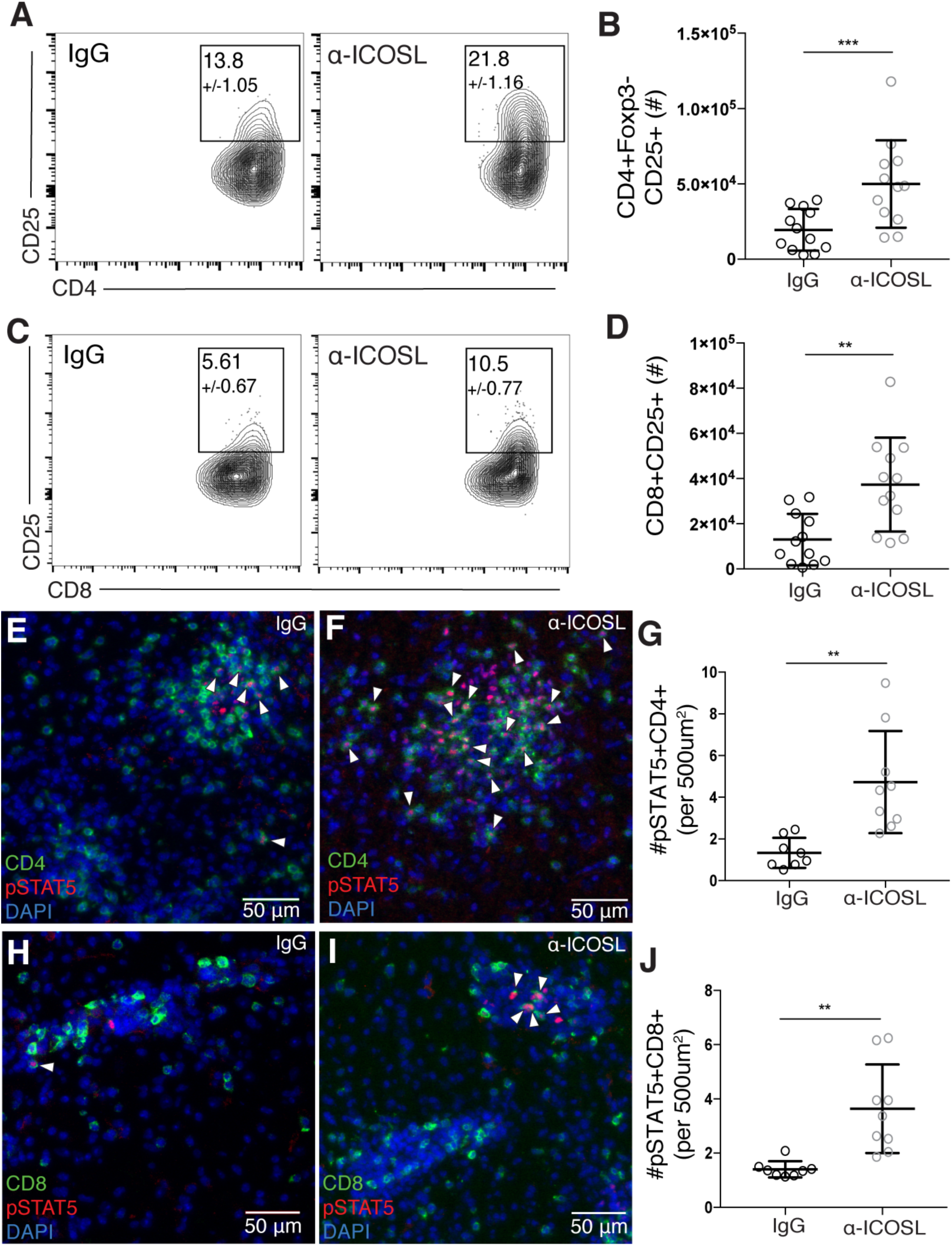
ICOSL blockade increases CD25 expression and STAT5 phosphorylation in effector T cells in the brain during chronic infection. (**A-D**) T cells were isolated from the brains of chronically infected control or α-ICOSL-treated mice. Representative flow plots of CD25+CD4+ effector T cells (**A**) and CD25+CD8+ effector T cells (**C**) are shown. Number in gate indicates the mean frequency of CD25+ cells +/- standard error. (**B**) Total number of CD25+ CD4+ effector T cells and (**D**) total number of CD25+ CD8+ T cells isolated from the brain (n=4 per group, data is pooled from 3 independent experiments and analyzed by randomized block ANOVA). (**E-I**) Brain sections from chronically infected control (**E, H**) and α-ICOSL-treated (**F, I**) mice were stained for CD4 or CD8 (green), pSTAT5 (red) and DAPI (blue). White arrowheads indicate pSTAT5+ CD4 or CD8 T cells. (**G**) The number of pSTAT5+CD4+ and (**J**) number of pSTAT5+CD8+ T cells were quantified per 500 µm^2^ (n=4-5 mice per group, data is pooled from two independent experiments and analyzed using randomized block ANOVA). * denotes p<0.05, ** denotes p<0.01, and *** denotes p<0.001 for all panels.

### IL-10R blockade does not affect Bcl-2 or CD25 expression on T cells in the brain during chronic infection

While increased expression of both Bcl-2 and CD25 was seen on effector T cells in the brain following ICOSL blockade, no change in Bcl-2 expression was seen in effector T cell populations isolated from the brains of α-IL-10R-treated mice (Figure 5A-B). Additionally, increased levels of CD25 were not observed to the same degree with α-IL-10R treatment, as there were no changes in the levels of CD25 expression on CD4+Foxp3-T cells, and only a slight increase in CD25 expression on the CD8+ T cell population (Figure 5C). Overall, though both IL-10R blockade and ICOSL blockade resulted in increased numbers of effector T cells in the brain, IL-10R blockade correlated with increased APC activation and inflammatory cytokine mRNA expression, while ICOSL blockade correlated with upregulation of IL-2-associated signaling molecules in effector T cells. These data further support that ICOS and IL-10 signaling pathways can differentially promote regulation of effector T cell responses in the brain during chronic infection.

**Figure 5.**
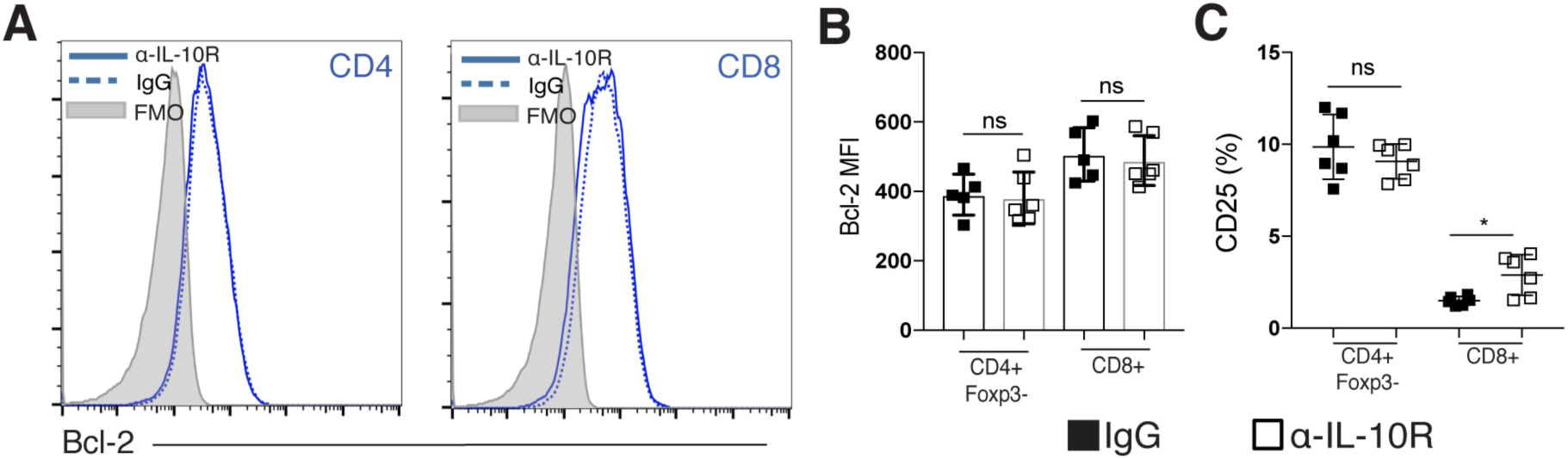
IL-10R blockade does not affect Bcl-2 or CD25 expression on T cells in the brain during chronic infection. (**A-C**) Effector T cell populations isolated from the brain were analyzed by flow cytometry following α-IL-10R blockade. (**A**) Representative histograms showing Bcl-2 expression on effector CD4+ and CD8+ T cells isolated from the brains of chronically infected mice treated with control or α-IL-10R blocking antibody. (**B**) The MFI of Bcl-2 on effector T cells isolated from the brains of control or α-IL-10R-treated mice (n=5-6 per group, data is representative of three independent experiments and analyzed using Student’s t test). (**C**) The frequency of CD25+ effector T cells isolated from the brains of chronically infected mice after α-IL-10R treatment (n= 6 per group, data is representative of two independent experiments and analyzed using Student’s t test). * denotes p<0.05, ** denotes p<0.01, and *** denotes p<0.001 for all panels.

## Discussion

Inflammation is required to promote clearance of pathogens, but preventing excessive inflammation during immune responses to infection is necessary to avoid immune-mediated tissue damage [3, 15, 57]. In cases of chronic infection, this balance between inflammation and regulation must be maintained over long periods of time [21, 43], but many of the signals required to maintain control of ongoing immune responses are not well understood. Our results identify one such signal, ICOS, whose expression on activated T cells in the chronically infected brain provides a suppressive signal preventing overabundant T cell accumulation in the inflamed CNS. We show that ICOSL blockade is associated with increased expression of IL-2-associated signaling molecules CD25, pSTAT5, and Bcl-2, as well as effector T cell accumulation in the brain.

ICOS, similar to its family member CD28, was initially characterized for its ability to amplify both B cell antibody production and T cell inflammatory cytokine production in cases of acute inflammation [24, 25, 53]. The pro-inflammatory role for ICOS signaling was subsequently supported by human data, as humans carrying mutations in the ICOS gene are included in the class of mutations known as common variable immunodeficiency (CVID) [58-60]. Patients with CVID have increased susceptibility to bacterial infections, and are diagnosed based on severely decreased class-switched antibody production, further implicating ICOS as important for T cell-dependent B cell responses [59]. Interestingly, further data from patients with CVID began to emerge highlighting widespread immune system abnormalities outside of the B cell compartment, particularly splenomegaly, a loss of naïve CD4+ T cells and expansion of activated CD4+ T cells, and increased T cell inflammatory cytokine production [58, 61, 62]. Additionally, despite being defined as an immunodeficiency disease, about 20% of CVID patients also present with autoimmune complications, though the pathogenesis of this autoimmunity remains unclear [63].

Studies regarding the role of ICOS in promoting regulation of immune responses largely come from mouse models, where ICOS has been shown to both support T_reg_ populations and promote IL-10 production [29, 31, 33, 34]. Surprisingly, we found no evidence of a local T_reg_ defect following ICOSL blockade, suggesting that T_regs_ in the brain during chronic infection rely on signals other than ICOS to support their survival and accumulation in the tissue, as well as their production of IL-10. Many activated effector T cells in the brain continue to produce IL-2 in the chronic phase of infection with *T. gondii*, so it is possible that the largely CD25+ T_regs_ in the CNS rely mainly on IL-2 signals for their maintenance in an inflamed tissue, whereas other signals are required during homeostasis when lower levels of IL-2 would be present in the absence of an ongoing effector T cell response

One of the main differences observed between IL-10R blockade and ICOSL blockade in the chronic phase of infection was the disparity in the magnitude of response. IL-10R is expressed more broadly than ICOS, which may explain the widespread inflammation and lethality of IL-10R versus ICOSL blockade. IL-10R is expressed by most hematopoietic cells, but can also be induced on non-hematopoietic cells such as fibroblasts and endothelial cells, rendering them also able to respond to IL-10 in inflammatory settings [64-66]. In the context of chronic *T. gondii* infection, this could explain why changes were seen in myeloid and T cell subsets in both the brain and peripheral tissues following α-IL-10R blockade. On the other hand, ICOS is only expressed on activated T cells during chronic *T. gondii* infection, and its highest levels of expression were found in the inflamed brain. This more limited expression pattern of ICOS compared to IL-10R could potentially explain the more local and specific response to ICOSL blockade.

Another interesting aspect of the immunological phenotype seen with IL-10R blockade during chronic infection was the differential effects on the accumulation of CD8+ and CD4+ effector T cell populations in the brain. Following IL-10R blockade, local APCs in the brain upregulate CD80, so it is possible that the CD4+ effector T cells infiltrating the brain are interacting with the highly activated local MHCII+ APCs more so than the CD8+ effector T cells, leading to their increased accumulation. It is interesting to note that, of the brain-infiltrating APCs isolated during chronic *T. gondii* infection, only a small fraction of them are classic CD8α+ cross-presenting DCs that could interact in a TCR-dependent fashion with infiltrating CD8+ T cells [67]. This could suggest that the activated CD8+ effector T cells in the brain rely less on local restimulation through TCR-MHC and costimulatory interactions, and are therefore less affected by extrinsic mechanisms of suppression through APCs in this context. Overall, it is largely unknown what kinds of secondary signals, such as TCR-MHC or costimulatory interaction, are required for activated T cells to carry out effector function and survive at a distal site of inflammation after initial priming in secondary lymphoid organs, though it is likely these requirements differ in some way for CD8+ and CD4+ T cells. Though the overall number of CD8+ effector T cells in the brain is not increased, we still observed an increased frequency of Ki67+ CD8+ T cells in the brain after IL-10R blockade, suggesting that this population is still responsive to IL-10-mediated suppression; perhaps through an intrinsic response to IL-10 that limits their proliferative capacity. IL-10 has been shown to be able to directly inhibit proliferation of CD8+ T cells *in vitro* [14], as well as control the threshold of their response to antigen upon initial activation [68]. Much is still unknown about the direct effects of IL-10 on activated CD8+ T cells that could contribute to their regulation, though these data further suggest that CD8+ effector T cells may be differentially regulated from the CD4+ effector T cell compartment in the brain.

Two well-characterized inhibitory co-receptors are CTLA-4 and PD-1, both of which have been shown to carry out their inhibitory effect at least partially through inhibition of the PI3K/Akt signaling pathway [69, 70]. In this light, it is interesting to consider ICOS, which is a potent activator of the PI3K/Akt pathway [71, 72], as also providing an inhibitory signal to T cells during chronic inflammation. Initial Akt activation induced during priming has been associated with increased T cell responses, both through promotion of T cell proliferation and survival [73-75]. However, more recent reports have shown that constitutive Akt activation in CD8+ T cells is associated with decreased expression of CD122 and Bcl-2, and can promote the development of short-lived effector T cells over the development of memory T cells, while constitutive STAT5 signaling can maintain Bcl-2 expression and favor the development of memory precursor cells [76, 77]. These results emphasize that the fate of T cells is extremely sensitive to both the level and duration of signaling cascades like PI3K/Akt. During the chronic neuroinflammation seen with *T. gondii* infection, continued activation of Akt downstream of ICOS in activated T cells in the brain could potentially serve as an intrinsic mechanism of controlling T cell responses by downregulating IL-2-associated signaling molecules and driving effector T cells in the brain to be short-lived effectors rather than memory precursors.

Overall, we provide evidence that ICOS costimulation provides an inhibitory signal to antigen-experienced effector T cells in the chronically inflamed brain. These data postulate a regulatory role for ICOS in T cells during chronic neuroinflammation that is distinct and more specific than the suppressive role of IL-10. Altogether, while we show that IL-10 is absolutely required for preventing exaggerated immune responses and immunopathology, other regulatory signals like ICOS are also likely at play during chronic inflammation that provide more fine-tuned suppression of ongoing immune responses without affecting IL-10-mediated suppression.

## Acknowledgements

The authors would like to thank all the members of the Harris lab and Center for Brain Immunology and Glia (BIG) for their insightful comments and discussion during the preparation of this work. We acknowledge support from the Research Histology core facility at the University of Virginia. We would also like to thank Marieke K. Jones for her helpful input regarding relevant statistical analysis and coding, and Dr. Ken Tung for his assistance with histopathological examination of tissue sections.

